# Disentangling the effects of climate change, landscape heterogeneity, and scale on phenological metrics

**DOI:** 10.1101/2021.02.05.429398

**Authors:** Erica A. Newman, Ian K. Breckheimer, Daniel S. Park

## Abstract

Phenology, the study of the timing of cyclical life history events and seasonal changes, is a fundamental aspect of how individual species, communities, and ecosystems will respond to climate change. Both biotic and abiotic phenological patterns are changing rapidly in response to changing seasonal temperatures and other climate-related drivers, and the consequences of these shifts for individual species and entire ecosystems are largely unknown. Landscape-scale simulations can address some of these needs for better predictions by demonstrating how phenology measures can vary with spatial and temporal grain of observations, and how phenological responses can vary with landscape heterogeneity and climate drivers. To explicitly examine the spatial and temporal scale-dependence of multiple phenology measures, we constructed simulated landscapes populated by virtual plant species with realistic phenologies and environmental sensitivities. This enabled us to examine phenology measures and environmental sensitivities along a continuum of spatial and temporal grains, while also controlling other aspects of sampling design. By relating measures of phenology calculated at a given spatiotemporal grain to average environmental conditions at that same grain size, we are able to determine observed environmental sensitivities for multiple phenological metrics at that spatial and temporal scale. We demonstrate that different phenological events change distinctly and predictably with spatial and temporal measurement scale, opening the way to incorporating scaling laws into predictions. Using plant flowering as our example, we identify that the timing of the beginnings or ends of an event (e.g., First Flower date, Last Flower date), can be especially sensitive to the spatial and temporal grain (or resolution) of observations. Our work provides an initial assessment of the role of observation scale in landscape phenology, and a general approach for incorporating scale-dependence into predictions of a variety of phenological time series.

## Introduction

Over the last two decades, the study of phenology, or the timing of biological and seasonal events, has taken on new relevance, as the effects of climate change have become increasingly noticeable. The consequences of phenological shifts and mismatches are unknown (Memmott et al. 2007). Much work has been done to assess the direction, magnitude, and mechanisms of phenological response across species, utilizing controlled experiments (Price and Waser 1998), remote sensing methods (X. Zhang et al. 2003), citizen science (Willis et al. 2017), natural history collections (Park et al. 2018), modeling, and combinations thereof. Phenological studies have long focused at the at the scale of individual organisms and plots, but large-scale digitization of natural history collections and survey data, as well as the advent of remotely sensed land surface phenology via satellite has increasingly facilitated research at more extensive taxonomic, spatial, and temporal scales over the last few decades. As a result, broad trends such as a general acceleration of plant phenology in response to warming, have emerged (Cleland et al. 2007).

However, a growing body of research suggests that phenological landscapes are highly complex, varying across spatial scales both within and among species (Körner and Basler 2010; Lapenis et al. 2014; Zohner and Renner 2014; H. Zhang et al. 2015; Cole and Sheldon 2017; Asam et al. 2018; Park et al. 2018). Though previous research spans diverse taxonomic, temporal, and spatial scales, harmonizing diverse scales of information has proven to be a challenge to the characterization of phenology, and we still lack a robust theoretical framework that can integrate this important body of knowledge (Newman et al. 2019; Gonzalez et al. 2020). Previous attempts to directly link observations made at different scales (e.g., ground-based observations of individuals vs satellite-derived landscape observations) have often yielded poor results (Chuine, Cambon, and Comtois 2000; Badeck et al. 2004; X. Zhang et al. 2017). Because scale and scaling are fundamental to ecological patterns including phenology (Woodcock and Strahler 1987; Levin 1992; Wiens 1989), synthesizing observations made at different spatiotemporal resolutions and extents are at the forefront of current phenological research (Cleland et al. 2007). Such efforts are necessary to provide accurate predictions about future global change impacts.

In this study, we use simulated datasets to compare spatiotemporal scaling across heterogeneous environments and demonstrate that the properties of phenological events can change predictably with scale. We thus provide a framework for increasing our understanding how the phenology functions at scales from the individual to the landscape via empirical and theoretical synthesis. In our instance, we use an empirically-informed simulation for virtual landscapes and species, to elucidate the inherent sensitivities of phenological metrics to measurement scale, independent of other factors, such as exogenous climate forcings. We demonstrate that the properties of multiple phenological events change distinctly from one another, but predictably with spatial and temporal measurement scale. Our simulation work highlights that some phenological measures (such as measures of peak or central tendency) are robust to large changes in spatial and temporal grain, while others are not. It also demonstrates that the effects of spatial and temporal sampling, aggregation, and scaling can be disentangled effectively from the effects of landscape heterogeneity and the effects of exogenous climate forcings on individual phenological metrics.

## Methods

### Landscape Phenology Simulations

Landscape simulation models have the potential to predict ecological metrics, including phenological time series, across scales, and provide a quantitative framework for investigating the implications of these predictions (Turner, Dale, and Gardner 1989; Wagner and Fortin 2005). To explicitly examine the spatial and temporal scale-dependence of multiple phenology measures, we constructed simulated landscapes populated by virtual plant species with realistic phenologies and environmental sensitivities. This simulation approach allowed us to examine phenology measures and environmental sensitivities along a continuum of spatial and temporal grains, while tightly controlling other aspects of sampling design. The simulated landscapes, and the species inhabiting them, were constructed to have similar properties to flowering plant communities in montane to subalpine environments in western North America, and were derived from a synthesis of plot-scale flowering phenology datasets across three locations: Mount Rainier National Park in the Washington Cascades (Theobald, Breckheimer, and HilleRisLambers 2017), MPG Ranch in the Sapphire Range of Montana (Durham et al. 2017), and Rocky Mountain Biological Laboratory in Western Colorado (Iler et al. 2017). As employed here, this method incorporates important phenological information and landscape heterogeneity factors for montane to subalpine environments in western North America, but could be used with other factors, simulated landscapes and species for other ecosystems.

To construct realistic phenological responses of virtual species, we first fit a hierarchical non-linear model describing species-specific phenologies and responses to climate for the combined three field datasets. Because the model drew species-specific parameters from statistical distributions, we could use this model to generate realistic phenological responses for 45 virtual species (Fig. 1). Virtual species are distinguished by their abundances, their means and variances for phenological response dates, and their peak abundance distributions across environmental gradients. Similarly, empirical measurements of microclimate at each field site were used to fit variogram models describing the pattern of spatial covariance of environmental variables at the study sites. These models were used to construct virtual landscapes with realistic spatial patterns of microclimate (Fig. 1), which in turn drive realistic spatial patterns of plant phenology. We then sampled plant phenology on the virtual landscapes at a variety of different spatial grains (from 2m −1024m) and temporal grains (sampling intervals from 1 −17 days), spanning the most prevalent spatial and temporal grains represented in the literature (Park et al., *in review*) (throughout the manuscript, spatial grains are reported as the linear measure of one side of a square unit, for example, 2m grain size corresponds to 4m^2^ area units). This allowed us to calculate a variety of measures of flowering phenology, including dates of first flowering, peak flowering, last flowering (or “First Flower,” “Peak Flower,” and “Last Flower,” respectively) and flowering duration, for each virtual species at each spatial and temporal grain.

**Figure 1.**
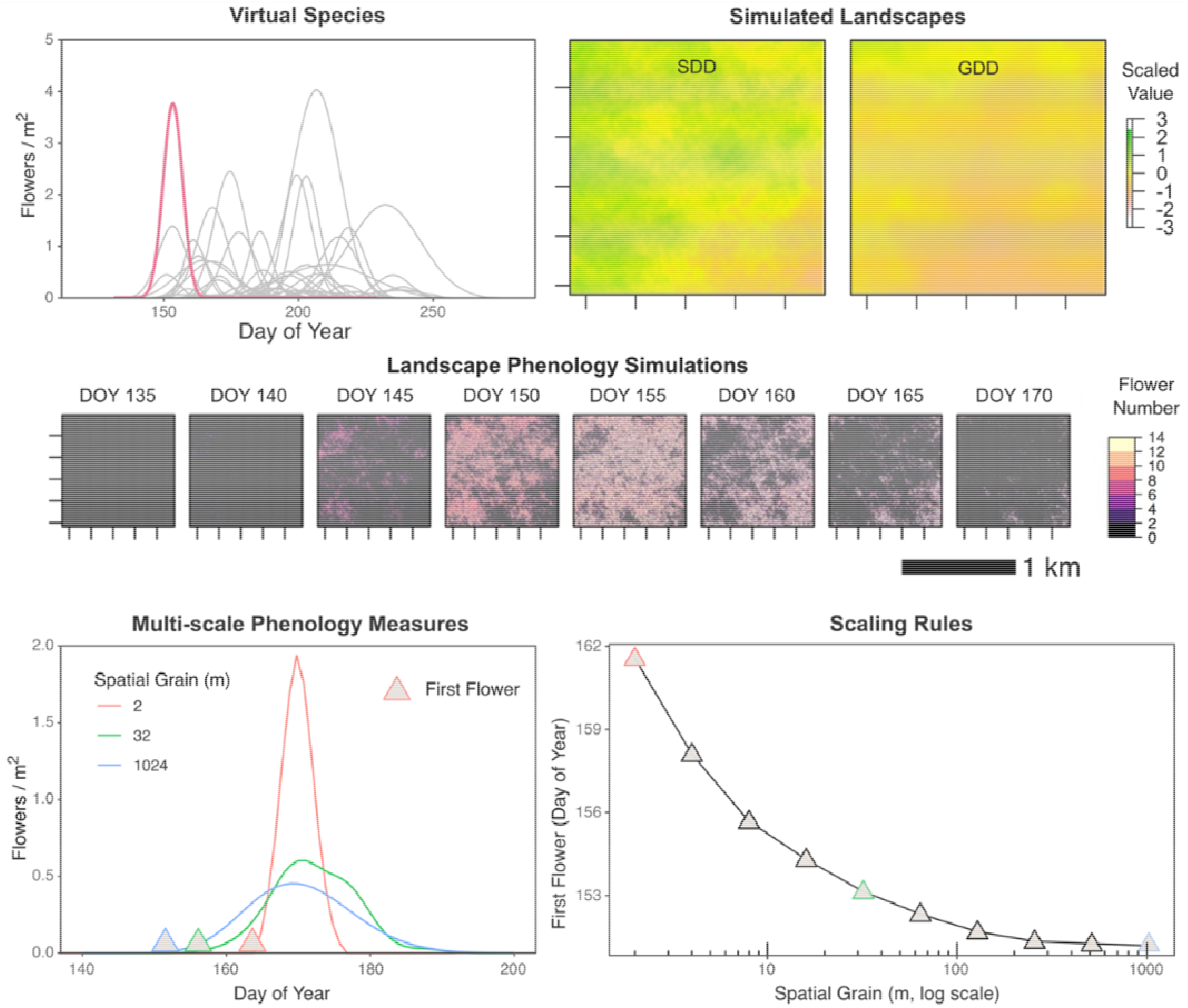
Summary of the simulation approach. Realistic flowering phenologies and responses to climate for virtual species (top-right panel), were generated from a Bayesian nonlinear model. Modeled flower densities were a function of day of year (DOY) and two climate variables: the snowpack disappearance day (SDD), and post-snow air temperature accumulation (Growing Degree-Days; GDD), both of which were allowed to vary across virtual landscapes as multivariate Gaussian random fields (top-right panels). Simulated flower counts were generated across the growing season on these landscapes (middle panels), and the progression of flowering was then summarized at a variety of spatial grains (bottom-left panel). Phenology measures such as date of First Flower were extracted from these time series at each spatial and temporal grain and used to examine scaling relationships (bottom-right panel).

Our approach makes use of “fully-nested” data structure, that is, data that have full spatial and temporal data associated with phenological events, which can be aggregated to increasingly coarser resolutions without loss of information. In the simulation, we have “perfect knowledge” of all phenological events, and from these, we can construct a scaling law related to what date of first, last, or peak event emerges from each resolution, up to the full spatial or temporal extent under consideration. This approach is similar to that sometimes used in macroecology, where mathematical scaling laws are constructed, and then extrapolated to unmeasured scales (Harte 2011; Harte and Newman 2014).For each phenology measure, we determined scaling effects by comparing the phenology measures computed at a given scale to the measures taken at the finest spatial and temporal scale available: 2m grain size and daily sampling. Code for the simulation and detailed methods can be found at: https://github.com/ibreckhe/phenoscaling_sims

Our simulation approach also allowed us to examine the scale-dependence of observed environmental sensitivities. The phenology of the virtual species respond to two aspects of the environment: the timing of seasonal snowpack disappearance (snow disappearance day, SDD) and the accumulation of air temperature forcing (growing degree-days) in the 90 days after snow disappearance (GDD), both of which vary across each virtual landscape. By relating measures of phenology calculated at a given spatial and temporal grain to average environmental conditions at that same grain size, we can determine observed environmental sensitivities at that spatial and temporal scale (Fig. 1). Because the “true” environmental sensitivities of these virtual species are known (as they were generated from distributions of sensitivities in the hierarchical model), we can measure scale effects by comparing the observed sensitivities at a given scale to the true values.

## Results

### Landscape phenology simulations

Simulated landscapes populated by virtual species can be used to examine the spatiotemporal scale-dependence of multiple phenology measures, and their sensitivity to environmental forcings. For instance, here, simulated landscapes and species were constructed to have similar properties to flowering plant communities in montane to subalpine environments in western North America, derived from a synthesis of plot-scale flowering phenology datasets (Theobald, Breckheimer, and HilleRisLambers 2017; Durham et al. 2017; Iler et al. 2017). Because the “true” environmental sensitivities of these virtual species are known, we were able to measure scale effects by comparing the observed sensitivities at a given spatial or temporal scale to their true values.

### Comparing the scale dependence of phenological metrics

To better understand the effects of scale and environmental forcings on phenological metrics, we investigated four measures of phenology – (the dates of) First Flower, Peak Flower, Last Flower, and Flowering Duration – and their sensitivity to environmental conditions using the simulation approach described in Figure 1. All examined metrics were scale-dependent, with some processes being more sensitive to statistical aggregation over time and space than others (Fig. 2). At coarser spatial grains, First Flower always appeared earlier, Last Flower always appeared later, and Flowering Duration therefore became longer. This was because coarser spatial samples incorporated more microclimate heterogeneity, and thus included some areas where flowering started earlier and later. At coarser temporal grains, observations of First Flower became later, Last Flower became earlier, and Flowering Duration therefore decreased. This was because less frequent observations were likely to miss the true start and end of the season, delivering estimates that skewed late for First Flower, and early for Last Flower. We consistently found that measures involving the start and end of the flowering season (Flowering Duration, First Flower, Last Flower), were considerably more scale-sensitive than the timing of Peak Flower, both in spatial and temporal grain.

**Figure 2.**
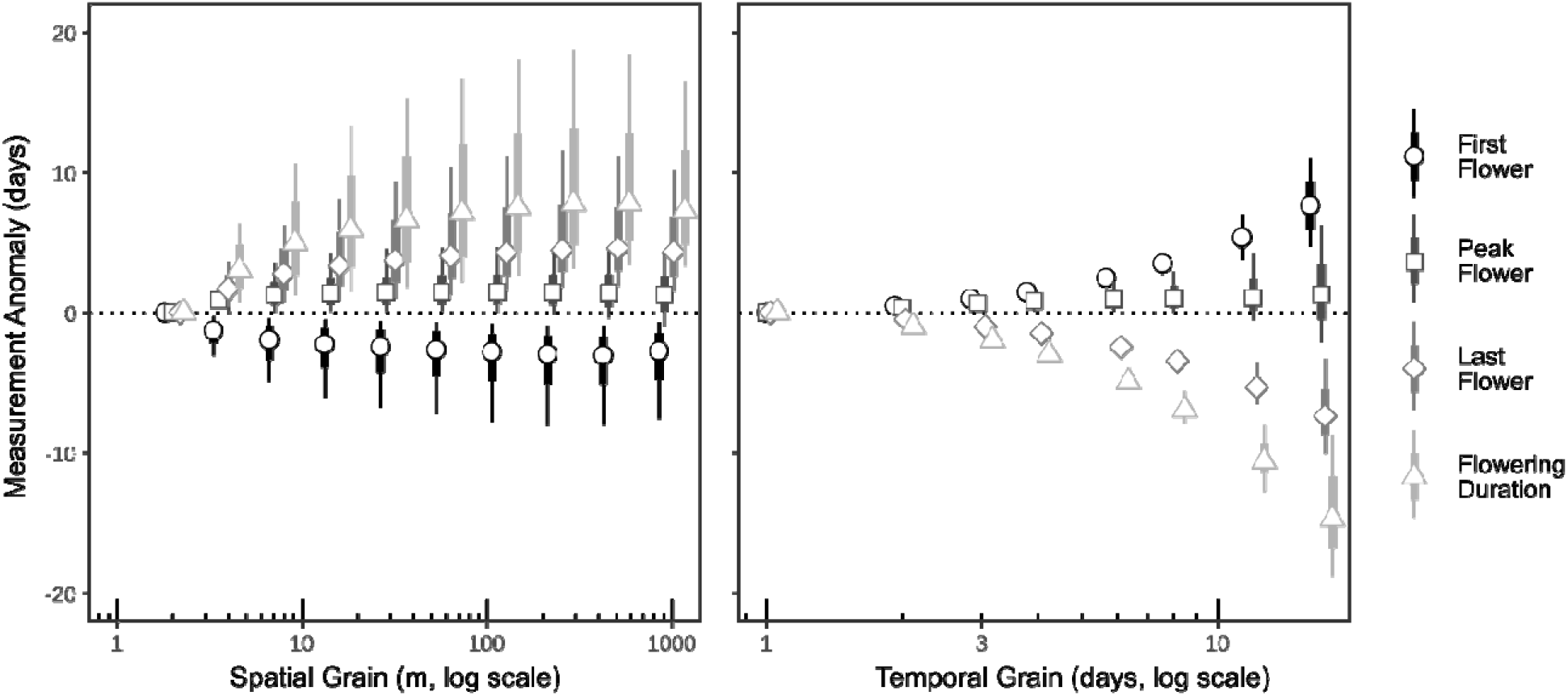
Simulation results demonstrating how phenology measures can vary with spatial and temporal grain of observations (left and right panels, respectively). Simulations place virtual plant species with a variety of realistic phenological responses to climate on virtual landscapes modeled after subalpine meadow ecosystems. Four phenology measures (First Flower, Peak Flower, Last Flower, and Flowering Duration) were computed after sampling these virtual landscapes at 10 different spatial grains (between 2 and 1000m), and 10 different temporal grains (between 1 per day and one per 17 days). Thick bars and thin bars represent 25/75% and 10/90% quantiles of the measures across all virtual species and landscapes.

### Scale dependence in phenological sensitivity

Observed phenological sensitivities can also be strongly scale-dependent (Fig.3, top panels). Spatial scaling effects caused sensitivity estimates to differ at 1km scales by up to +0.38 days per snow disappearance day, and by up to 0.02 days per accumulated °C compared to estimates at the finest spatial grain of 2m (Fig.3, top panels). The environmental sensitivities of start and end of season measures were considerably more scale dependent than the environmental sensitivity of Peak Flower, which was essentially stable across the spatial grains we tested. For most of the virtual species and landscape combinations, changes in spatial grain altered the magnitude, but not direction, of expected phenological shifts in response to changing forcing. For some species/landscape combinations, however, shifts in observation grain caused environment – phenology relationships to change sign. This was especially common for the environmental sensitivities of Flowering Duration, which changed sign in 24% of species/landscape combinations for SDD, and 21% of combinations for GDD at 1km spatial grains, compared to 2m grains (Fig. 3, bottom panels). These results highlight the importance of spatiotemporal grain in the reporting and analysis of phenology measures, especially for those that correspond to the start or end of a process.

**Figure 3.**
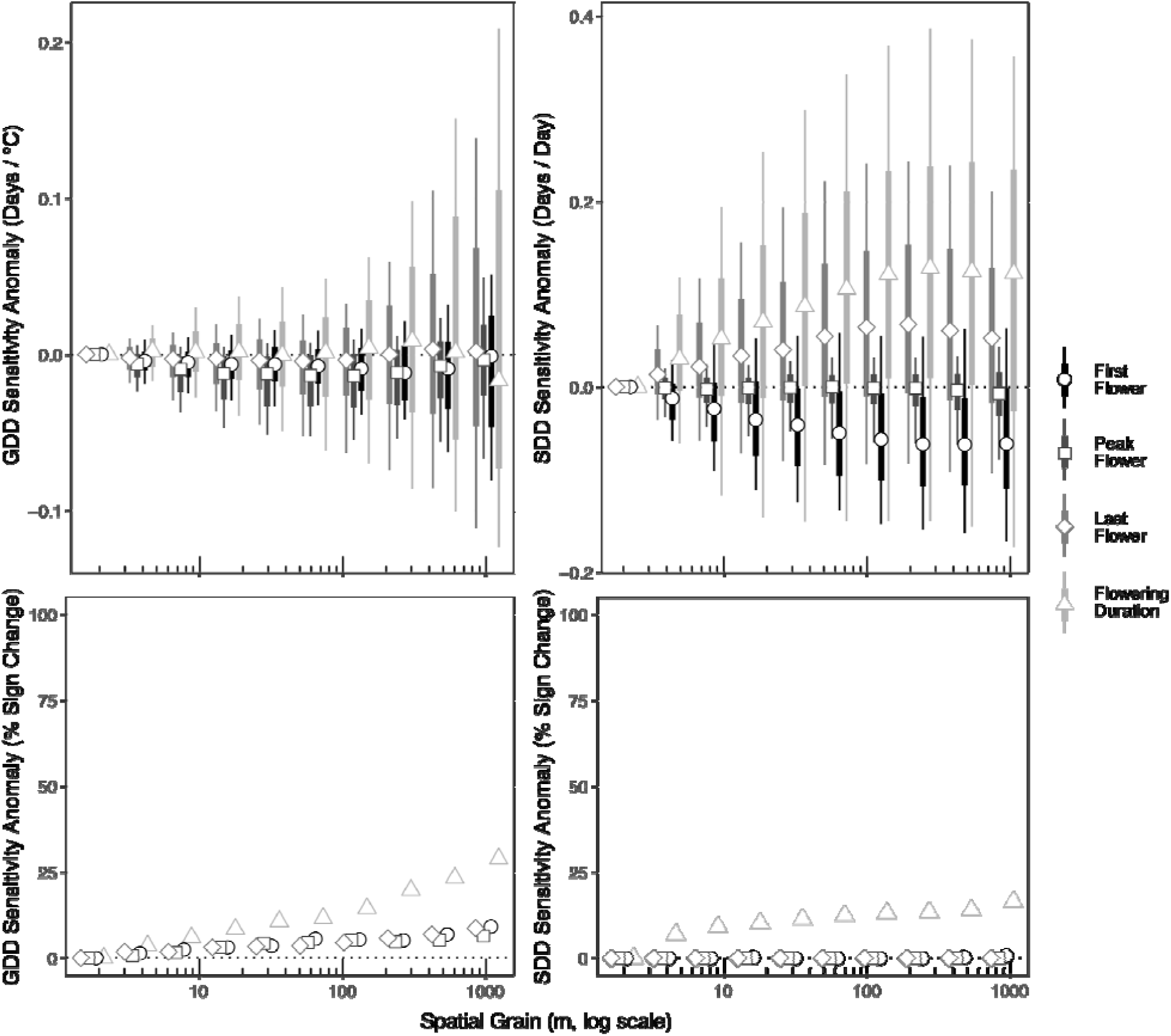
Simulation results demonstrating how environmental sensitivities can vary with the spatial grain of observations. Climate variables driving plant phenology were spring air temperature (Growing Degree-Days or GDD; left panels), and the timing of snowpack disappearance (SDD; right panels). Top panels show the change in environmental sensitivities as a function of spatial grain. Thick bars and thin bars represent 25/75% and 10/90% quantiles of the measures across all virtual species and landscapes. To put these anomalies in proportion, the median GDD sensitivity across all species and phenology measures at 2m grain was 0.10 days / 10 °C, and the median SDD sensitivity was 0.52 days / day. Bottom panels show the percent of virtual species and landscapes where observed environmental sensitivities at a given scale were of a different sign than sensitivities at the finest spatial grain (2m).

## Discussion

Understanding and predicting the timing of phenological events is critically important to ecologists, conservation biologists, and evolutionary biologists. Climate change simultaneously alters multiple ecological axes, and phenological events are among the most prominently affected (Wolkovich, Cook, and Davies 2014). The timing and location of phenological events provides structure to plant communities and their associated mutualists and predators, and although consequences of disruptions to these patterns have unknown consequences, lack of availability of resources at critical times are expected to negatively impact abundance of individual species as well as community structure, and may lead to extinctions (Memmott et al. 2007). Spatiotemporal biodiversity increases niche complementarity in species interactions and affects resource partitioning, reducing competition among co-occurring species (Venjakob et al. 2016). Thus, changes in temporal plant community composition can affect resource availability, trophic interactions, diversity of associated animal communities, and ecosystem services (Corlett and Lafrankie 1998; Edwards and Richardson 2004; Post and Forchhammer 2008; Sackett et al. 2011; Kudo and Ida 2013; Kendrick et al. 2015). Despite the importance of the interaction between climate and phenology, we have lacked an understanding of key scale-dependent mechanisms that influence phenological responses across landscapes. Indeed, it has become increasingly clear that we cannot simply extrapolate phenological knowledge across scales (Tian et al. 2020; Xie and Wilson 2020).

The scale dependence we observe in the phenological responses of species and communities can be attributed to a number of factors. These include environmental heterogeneity, variation in species’ sensitivities to environmental forcings across the landscape, as well as artifacts of statistical aggregation (Levin 1992). To account for these effects when integrating knowledge across scales, it is necessary to not only quantify the degree of scale dependence, but to elucidate the cause. The simulation approach we outline provides a way to address this issue and facilitate the informative integration of phenological information across scales.

We recognize several limitations to our study, enumerated here: (1) Our simulation approach assumes that the same level of detail is captured at every scale of observation; (2) Our simulated landscape and species were based on empirical data on flowering plant communities in montane to subalpine environments in western North America; (3) The results of our simulations may not apply universally across systems and all measures of phenological events, however, they should be general enough to apply to systems with well-defined seasons, and spatial domains; (4) The type of data needed to extract these scaling laws have the requirement that they be “fully-nested,” that is, to have full spatial and temporal data associated with phenological events that can then be aggregated to coarser and coarser resolution. However, recent advances in remote sensing technologies and machine learning applications are making it increasingly possible to overcome these limitations, and to identify functional types, species, and even individuals from large scale data collected from phenocams, drones, and satellites (Assmann et al. 2020; Rossi et al. 2019). Furthermore, our simulation approach can be adapted to less than ideal datasets to parse and account for at least a portion of the variation in phenological measurements among studies conducted at different scales.

We present a conceptual framework for landscape-scale simulations of phenological time series that builds off of multiscale observations to better investigate how the seasonality of ecosystems across landscapes and seascapes respond to environmental variability and change. Along these lines, we provide an example approach for estimating scale dependence for phenological metrics for plants across both spatial and temporal grain and resolution, which makes use of fully-nested data structures. We provide guidance on how to create null models for spatial and temporal scaling with individual phenological metrics, as well as software code in support of these null models. These methods can be easily adapted to other phenological metrics, landscapes, and ecosystems. These efforts may lead to better disentangling of the effects of landscape heterogeneity and scale from the true effects of climate change. We thus set the stage for a new generation of empirical research in the field that builds off of multi-scale observations to understand how phenology across Earth’s ecosystems respond to environmental variability and change.

## Acknowledgements

We thank DH Hembry for manuscript edits and discussions. We also thank BJ Enquist and members of the Bridging Biodiversity and Conservation Science (BBCS) program at the University of Arizona for their comments on earlier versions of this work. DSP was supported by NSF-DEB 1754584, IKB was supported by a National Science Foundation Postdoctoral Fellowship in Biology (PRFB-1711936).

